# A phylogenetic model for the recruitment of species into microbial communities and application to studies of the human microbiome

**DOI:** 10.1101/685644

**Authors:** John L. Darcy, Alex D. Washburne, Michael S. Robeson, Tiffany Prest, Steven K. Schmidt, Catherine A. Lozupone

## Abstract

Understanding when and why new species are recruited into microbial communities is a formidable problem with implications for managing microbial systems, for instance by helping us better understand whether a probiotic or pathogen would be expected to colonize a human microbiome. Much theory in microbial temporal dynamics is focused on how phylogenetic relationships between microbes impact the order in which those microbes are recruited; for example species that are closely related may competitively exclude each other. However, several recent human microbiome studies have observed closely-related bacteria being recruited into microbial communities in short succession, suggesting that microbial community assembly is historically contingent, but competitive exclusion of close relatives may not be important. To address this, we developed a mathematical model that describes the order in which new species are detected in microbial communities over time within a phylogenetic framework. We use our model to test three hypothetical assembly modes: underdispersion (species recruitment is more likely if a close relative was previously detected), overdispersion (recruitment is more likely if a close relative has not been previously detected), and the neutral model (recruitment likelihood is not related to phylogenetic relationships among species). We applied our model to longitudinal human microbiome data, and found that for the individuals we analyzed, the human microbiome generally follows the underdispersion (i.e. nepotism) hypothesis. Exceptions were oral communities and the fecal communities of two infants that had undergone heavy antibiotic treatment. None of the data sets we analyzed showed statistically significant phylogenetic overdispersion.

## Introduction

A central question in microbial community assembly, especially in the human microbiome, is when and why microbes are recruited into communities. If there are patterns or general rules for which taxa have higher probabilities of recruitment, these rules can guide habitat restoration projects, help us better understand whether probiotics or pathogens will colonize, and better exploit disturbance as a tool for managing microbial systems related to human health and disease. Recruitment of new species can be studied by evaluating the order in which species are empirically detected in time-series experiments, given data such as which species have already been detected or what changes occur in an environment over time [1; 2]. Although a changing environment clearly selects for new species, it has also been shown that microbial community structure is often historically contingent on previous states of that community [1; 3; 2; 4; 5]. This reflects not only that microbial communities are temporally autocorrelated (gradual change over time), but also that the recruitment of a given species is a function of which species in the community are already present or have modified the local environment. Such historically contingent patterns have mainly been observed and tested within a phylogenetic context, because amplicon data naturally lend themselves to the creation of phylogenies, and because phylogenies have been shown to be predictive of genomic (and perhaps niche) overlap in human associated microbiota [6; 7] and in general [8; 9].

Within this phylogenetic framework, a predominant hypothesis has been that closely related microbes inhibit each other’s successful recruitment [1; 3; 4]. The proposed mechanism for this hypothesis is that closely related microbes likely have similar niches (phylogenetic niche conservatism [10; 8; 9]), and species already established within a community will occupy their niches to the exclusion of ecologically similar organisms. This is also the basis of Darwin’s naturalization hypothesis [11], which proposed that new species are less likely to be recruited if a close relative is present [12]. Indeed, this assembly mode has been found to be the case in artificial nectar microcosms, where phylogenetically similar yeast species had similar nutrient requirements, and inhibited each others’ colonization [13]. If competitive exclusion of closely-related species is the predominant mode of microbial community assembly in the human microbiome, close relatives would be less likely to be recruited into the same community, as compared to more distant relatives. In this paper, we refer to assembly where distant relatives are more likely to be recruited into a community than close relatives as the **overdispersion hypothesis**, since it predicts the preferential addition of novel phylogenetic diversity to a community (*i.e.* phylogenetic overdispersion).

Overdispersion is far from universal, and multiple studies have shown that extremely close relatives can coexist within the human microbiome [14; 15; 16], and may even be preferentially recruited [17]. This is consistent with simulations showing that clusters of closely-related species can persist despite strong within-cluster competition, when immigration rate is high [18]. Indeed, Darwin’s pre-adaptation hypothesis predicts that species with a close relative present in a community will be preferentially recruited, because they are likely to already be adapted to the new environment [11]. Also, more recent ecological theory posits competition between distantly-related species may result in phylogenetically clustered communities, although it may be more appropriate to include this type of competition within selection [19]. Per the empirical observations and reasoning above, we might expect that new close relatives are more likely to be detected than new distant relatives, so the amount of new phylogenetic diversity added to a community is minimized (phylogenetic underdispersion). Thus, for empirical data our **underdispersion hypothesis** is that species are more likely to be detected when they have a close relative that was previously detected, predicting preferential addition of minimal novel phylogenetic diversity (phylogenetic underdispersion). The over- and underdispersion hypotheses are alternatives to the null hypothesis that recruitment is independent of phylogenetic relatedness among species. Since the null hypothesis is species-neutral (and phylogenetically neutral), we refer to it as the **neutral hypothesis**.

It should be noted that our use of the terms “overdispersion” and “underdispersion” are slightly different in this manuscript compared to use of the same terms elsewhere. In many cases, these words refer to the state of a community at a single timepoint or sample, with overdispersion indicating more diversity in that sample than expected by chance, and underdispersion indicating less [20]. Instead, our use of over- and underdispersion refers to the amount of newly added diversity over time. In our overdispersion hypothesis, phylogenetically novel species are preferentially added to communities, meaning more new diversity is added than expected by chance. Under our underdispersion hypothesis, the reverse is true. Following this, our question concerns the order in which new species are recruited in a time-series, rather than community composition of any given sample. Furthermore, while we are interested in the biological phenomenon of species recruitment, the empirical result of recruitment is detection. Evidence of recruitment is a lack of detection, and then subsequent detection of a species via high-throughput DNA sequencing data. It is possible for a species to have been recruited into a community but not be detected, although this source of experimental error diminishes as sequencing depth increases. Also, the extent to which a species has actually been recruited into a community is questionable, if it is sufficiently rare that it is not detected in an Illumina sequencing run with tens of thousands of reads per sample (e.g. [21]). Still, we have taken care to use the term “recruitment” when discussing the biological phenomenon under investigation, and “detection” when describing empirical data and results.

Here, we use the phylogenetic relationships among species within a time-series to test the extent to which our over- or underdispersion hypotheses hold true. Instead of analyzing broad patterns of community change via beta-diversity statistics (*e.g.* UniFrac [22]) or analyzing patterns of select clades within the community (*e.g.* PhyloFactor [23], Edge PCA [24]), we model the probability of new species’ recruitment into a community for the first time as a monotonic function of their phylogenetic distances to members of the community that have already been recruited.

The model we present here can be used to estimate the degree to which the recruitment of new species is more or less likely when a close relative has been previously recruited. We fit our model to several time-series human microbiome data sets [25; 26; 21] to compare the strength of under- or overdispersion between subjects, sample sites, or time periods. We found that for the data sets we analyzed (36 individuals across 3 studies), the human microbiome generally follows the underdispersion hypothesis. There were exceptions where this pattern was not significantly different than the neutral model, but none of the longitudinal data sets we analyzed showed statistically significant overdispersion.

## Materials and Methods

### Overview

With our model, our goal is to estimate the extent to which recruitment of new species over time is related to the new species’ phylogenetic similarity to (or distance from) species that were already recruited at previous timepoints. Our **Statistical Model** describes the probabilities of detecting new species over time. We use our model with empirical data via **Simulations**, where we re-sample the empirically detected species using our model with known parameter values, to produce surrogate data sets. Specifically, we fix and record the model’s dispersion parameter (*D*), which determines the extent to which species with a close relative are preferentially added to the surrogate community (or, conversely, if species without a close relative are preferred). Our **Parameter Estimation** compares the empirical pattern of species detection to that of the surrogate data sets (which have known *D* values), in order to determine which value of *D* best describes the empirical data. **Hypothesis Testing** is done by comparing empirical data to repeated simulations under the neutral model, which is *D* = 0. We describe the bioinformatic and technical details of this process in our **Analysis** section, and make our code available to others in the **Code and Data** section.

### Statistical Model

At any point in time, a community is composed of many species, and other species are not present but are available to be added (“species pool”). Our model parameterizes the probability of detecting species in a local community for the first time, based on their phylogenetic distances from species that have already been detected. In a species-neutral model of community assembly, each species *i* in the species pool has the same probability of detection at time *t*, irrespective of how different it is from species that have already been detected. Thus, the neutral model for first-time species detections is a random draw without replacement of species from the species pool. We extend the species-neutral model by modeling the probability *p*_*it*_ of species *i* being detected for the first time at time *t* as,

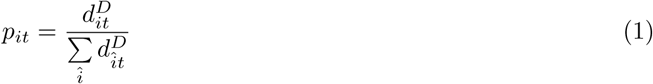

where *d*_*it*_ is the phylogenetic distance from species *i* to its closest relative that has already been detected prior to timepoint *t*, and *D* is a dispersion parameter.

When *D* = 0, our model functions as a neutral model; all species have the same probability of being detected for the first time, since *p*_*it*_ is the same for every species. When *D* < 0, *p*_*it*_ decreases with *d*_*it*_ meaning that species from the species pool have higher probabilities of detection when they are more closely related to species that have already been detected in the local community (underdispersion; phylogenetically constrained). When *D* > 0, the opposite is true (overdispersion; phylogenetically divergent). Our hypothesis testing and parameter estimation focus on the dispersion parameter, *D*.

### Simulations

Our analysis of a data set relies on re-constructing that data set via simulation of our statistical model using known values of 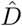, allowing for hypothesis testing and parameter estimation (we refer to the empirical dispersion parameter as *D*, and use 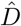 to refer to surrogate values used in simulations). Using the empirical data as a starting point, we simulate many surrogate data sets with 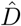 values ranging from 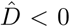 (under-dispersed) to 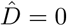 (neutral) to 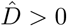 (overdispersed). This is done so that the empirical data can later be compared to the surrogate data sets, to estimate the empirical value of *D*.

We start each surrogate data set with the same species present in the first sample in the time-series of its corresponding empirical data set. Then, surrogate data sets are constructed forward in time by randomly drawing *r*_*t*_ new species from the species pool, where the probabilities of detecting those species are given by Equation 1, and *r*_*t*_ is the number of new species detected in the empirical data set from times *t* − 1 to *t*. The number of new species detected from the empirical data set is used so that species richness is kept constant between the empirical data set and all surrogate data sets. The species pool is updated to exclude those species drawn at previous timepoints, and the newly sampled species are recorded. Surrogate data sets are produced for many different 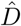 values, ranging from underdispersed to overdispersed models. We performed 500 simulations (as described above) for each data set analyzed.

### Parameter Estimation

Our main goal is to estimate the empirical dispersion parameter *D* (Equation 1), which quantifies the degree to which first-time species detections are phylogenetically underdispersed (*D* < 0), neutral (*D* = 0), or overdispersed (*D* > 0), corresponding to our hypotheses. To this end, we use Faith’s phylodiversity [27] to compare each of the 500 surrogate data sets (described above) to the empirical data set. Phylodiversity is the sum of branch-lengths on a phylogenetic tree for a set of species, so phylodiversity of a set of highly related species is low (phylogenetically constrained) because there are no long branch lengths in the tree, but phylodiversity is higher (phylogenetically divergent) for a set of more distantly related species [27]. If *D* ≠ 0, then species are preferentially added if they have relatively low (*D* < 0) or relatively high (*D* > 0) phylogenetic distance to the resident community (*d*_*it*_, Equation 1), yielding accumulations of total phylodiversity that are relatively slow (*D* < 0) or relatively fast (*D* > 0) compared to the neutral model (Fig. 1A). In other words, at any timepoint *t*, the phylogenetic diversity of species that have already been observed is *PD*_*t*_, and the extent to which *PD*_*t*_ accelerates or decelerates over a sampling effort depends on *D*. Because of this, we can estimate *D* by comparing the empirical phylodiversity curve to our surrogate phylodiversity curves, which have known 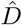 values.

**Fig. 1.**
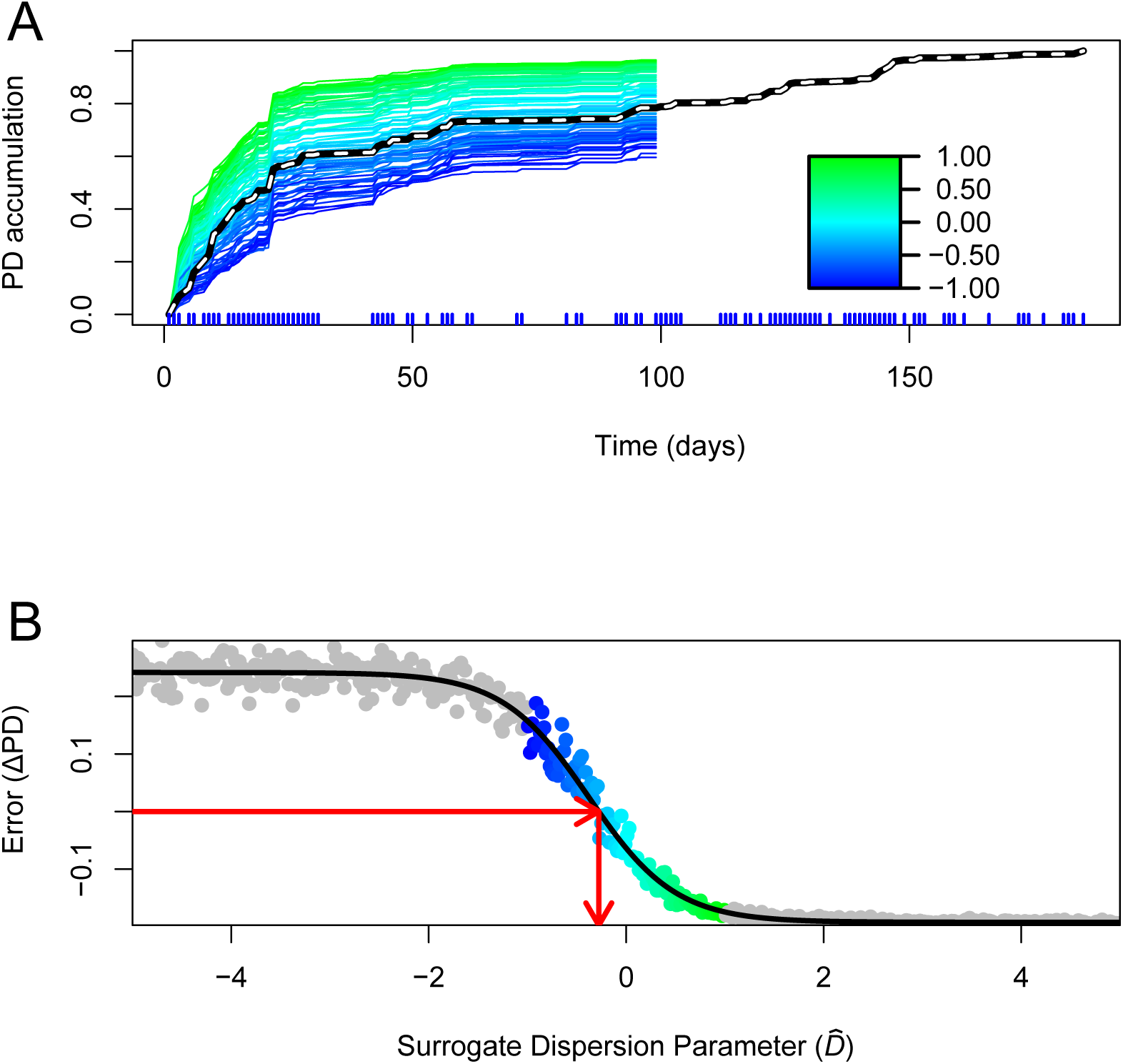
Phylodiversity accumulation and model fitting in the female feces data set [25]. Plot A shows empirical (dashed) and surrogate phylodiversity accumulation curves. Surrogate curves are colored according to 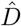 value (Equation 1). New species that have a previously-detected close relative contribute little phylodiversity and cause slow phylodiversity accumulation (blue). New species that do not have a close relative contribute more phylodiversity and cause faster accumulation (green). The empirical model (dashed) is below the neutral model (teal), signifying underdispersion in the order of first-time species detections. The times of sampling points are shown as vertical blue lines below the X-axis. Curves are rescaled from 0 to 1 in this figure. Plot B shows how empirical and surrogate data are compared to generate an estimate for *D*. Differences between empirical and surrogate data at time *m* are shown on the Y-axis, and the 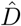 values used to generate surrogate data sets are shown on the X-axis. Color-coded points correspond to surrogate data sets shown in plot A. Values shown in gray result from using extreme values of 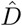, which help the logistic error model (black line) fit to the data, and are not shown in plot A. The red arrows show the process of error minimization, yielding a *D* estimate. A figure showing significance testing for these data is available as Fig. S1.

For the comparison of an empirical phylodiversity accumulation curve to curves for corresponding surrogate data sets, we evaluate the amount of phylodiversity *PD*_*m*_ accumulated at time index *m*, midpoint between the first and final samples. Time *m* is used because this leaves many species yet to be observed in the species pool, so that there can be variability in surrogate data sets. Multiple time indices are not used to compare surrogate and empirical data sets because each value 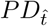 is a function of all values 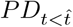. *PD*_*m*_ values are calculated for all surrogate data sets, and a *PD*_*m*_ value is calculated for the empirical data set. The difference between the empirical *PD*_*m*_ and *PD*_*m*_ simulated with 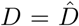 is 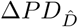, which is the error between surrogate and empirical data. We then estimate the empirical value of *D* by minimizing 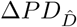 (Fig. 1B). This minimization is performed using a logistic error model,

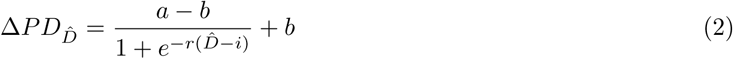

where *a* and *b* are the upper and lower horizontal asymptotes, and *r* and *i* are rate and inflection parameters for the logistic model. 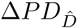 is modeled with a logistic function because there is a maximum and minimum observable 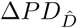 value as a function of the phylogeny; this is because there are strict minimum and maximum limits to the amount of phylodiversity obtainable by observing *n* species where *n* is the total species richness accumulated up to time *m*. The two horizontal asymptotes of the logistic model are easily fit to these extremes (Fig. 1B). Once fit, the error model is solved for Δ*PD* = 0, giving an estimate for the empirical *D*. Confidence intervals for this estimate are obtained via bootstrapping our error model.

### Hypothesis Testing

For this test, our null hypothesis is the neutral model, where *D* = 0, since this model represents the absence of the effect we are testing. We test this null hypothesis competitively by simulating 1000 surrogate data sets at *D* = 0 (Fig. S1A) to generate a null *PD*_*m*_ distribution. The empirical *PD*_*m*_ is compared to this distribution (Fig. S1B), and if the empirical *PD*_*m*_ is below the 2.5% quantile or above the 97.5% quantile, we reject the null (*i.e.* neutral) hypothesis. Evidence of either overdispersion (*D* > 0) or underdispersion (*D* < 0) allows us to reject. A second method of simulating communities under D = 0 was used as well, where simulated communities were drawn without replacement from the pool of all individual observations within a rarefied data set. In this context, an “observation” is a datum within a species-by-sample matrix of count data; a given column contains d observations where d is the sequencing depth. This “individual null” accounts for differences in total species relative abundance across a time series, while the simulation above only considers presence-absence of species. Results of this model were computed the same as above.

### Analysis

This section is a summary of our data analysis. Detailed methods for this section are available as supplemental information.

We ran our model on data from 36 individuals from three data sources. Two individuals were from Caporaso *et al.* [25], 33 were from Yassour *et al.* [21], and one was from Koenig *et al.* [26]. In all cases, data were downloaded and processed using the unoise3 pipeline [28], which clusters sequence data into exact sequence variants called zOTUs. The Koenig *et al.* infant gut data set was split into two data sets, one for samples collected before the subject began consuming baby formula, and one after. Our model was run on these data as described above, resulting in *D* estimates for the before and after formula data sets.

The “moving pictures” [25] data were split into eight data sets, one for each combination of subject (n=2) and body site (feces, right and left palms, tongue), and our model was run on each of these data sets. Analyses of these data was also done using two approaches that allowed us to test the importance of the set of species that are included in the species pool. One alternate approach analyzed communities in a “meta” context, where the species pool for a given palm was composed of all four palms in the whole data set. If we were to estimate similar *D* values for both the “meta” and “self” analyses, the inclusion of extra species in the species pool would be of little importance to the model. The other alternate approach analyzed data using a sliding-window approach, wherein our model was run separately on multiple overlapping windows of 5 consecutive days within the same data set, in order to see how *D* varied over time.

Finnish infant sequence data from Yassour *et al.* [21] were split into data sets for each of 33 individuals, and our model was run for each. Estimated *D* values were compared between subjects that had been treated with oral antibiotics (n=18) and subjects that had not (n=15) using a Mann-Whitney test. Because this data source had so many subjects, we used these data to test whether the number of zOTUs, total phylodiversity, or number of timepoints had an effect on *D* estimates via correlation analysis.

### Code and Data

R code and data to replicate our analysis, or to perform a similar analysis on other data, are available on GitHub, at https://github.com/darcyj/pd_model.

## Results

By varying 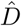, we successfully changed the rate at which phylodiversity is added to surrogate (*i.e.* resampled) microbial communities over time (Fig. 1A). Compared to the neutral model where 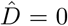, higher 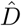 values result in phylodiversity accumulating quickly, since in the overdispersed model, species that contribute more phylodiversity are preferentially sampled. Conversely, lower 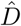 values result in phylodiversity accumulating slowly, since in the underdispersed model, species that contribute less phylodiversity (since they are very similar to species that are already present) are preferentially sampled. These results show that the *D* parameter in our model successfully corresponds to over- and underdispersion relative to the neutral model. Our error model also fit well to the differences between empirical and surrogate data sets (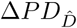, Fig. 1B). Each error model fit was visually inspected for goodness of fit, to be sure that *D* estimates were not spurious. All data sets passed this inspection.

### Results from “moving pictures” data

All time-series from adult feces and palm microbiomes [25] showed significant phylogenetic underdispersion of first-time zOTU detections (Fig. 2). This means that when a zOTU was detected for the first time in one of these communities, it was more likely to be phylogenetically similar to a zOTU that had previously been detected in community. For both the male and female subject, *D* estimates were lower (more underdispersed) in the feces than in the palms, left and right palm *D* estimates were similar to each other, and tongue *D* estimates were higher. All sites except the tongue showed statistically significant underdispersion in both subjects, while tongue data were not significantly different than the neutral model. The results for the “individual null” were the same. In the comparison between “meta” and “self” models, “meta” models needed to be much more underdispersed than “self” in order to approximate empirical phylogenetic diversity accumulation (Fig. S2). We also observed a general upward trend in *D* in our sliding window analysis of the male right palm data set (Fig. S3), although this trend was only observed over 19 days.

**Fig. 2.**
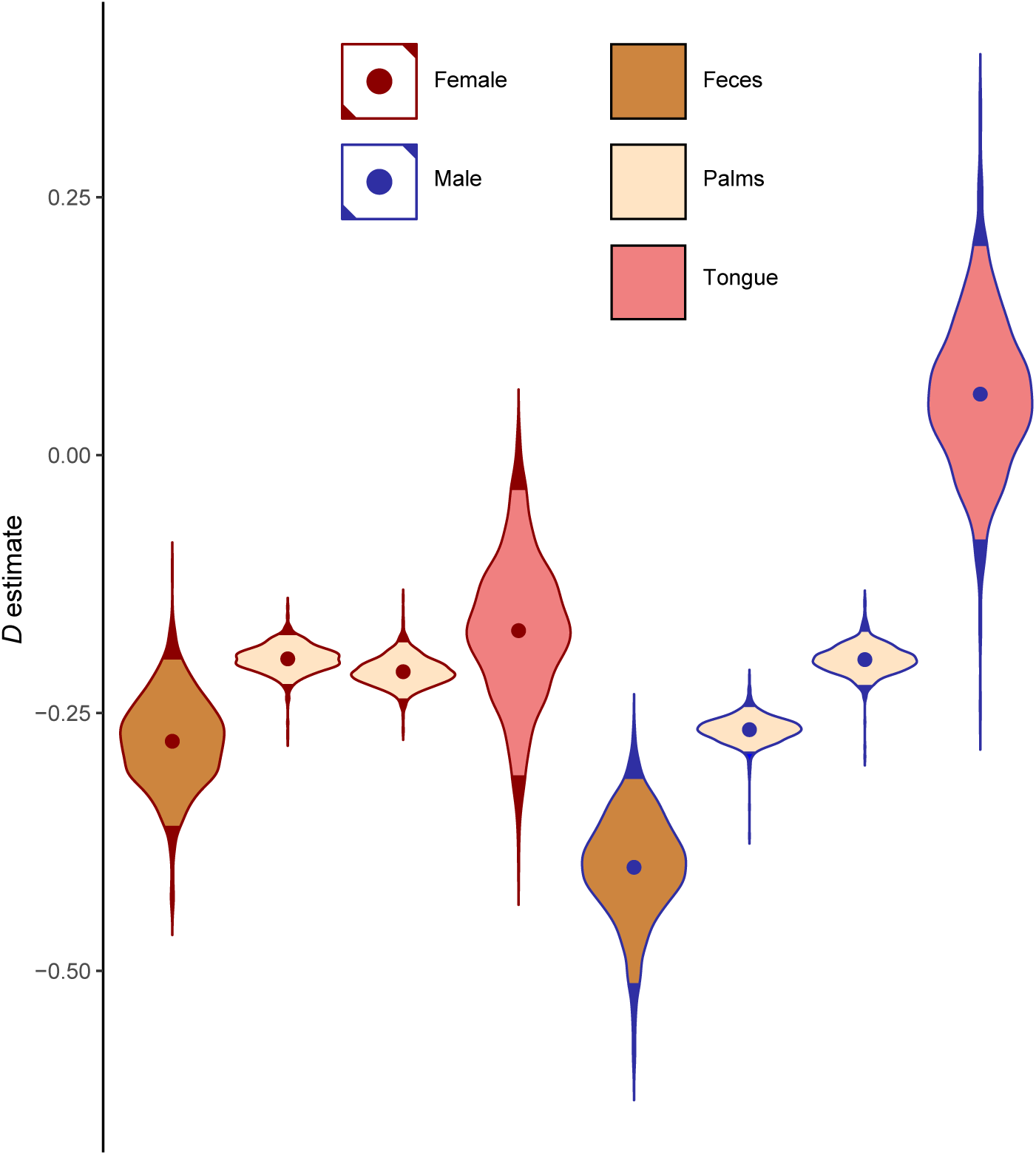
Dispersion parameter (*D*) estimates for “moving pictures” [25] data sets. The subject’s sex is shown as the outline color of each violin, and the body site is shown as fill color. The four body sites for the female subject are shown at left, and the four body sites for the male subject are shown at right. Each violin shows the distribution of *D* estimates given by logistic error model bootstraps, and the dots within violins are means. Light-colored portions of violins represent 95% of bootstraps. The two subjects analyzed show parallel *D* estimates, with feces being the lowest, followed by palms which are all similar, followed by tongue communities. For both subjects, tongue patterns were not significantly different than the neutral model.

### Results from infant gut data

Empirical phylodiversity accumulation in the infant gut microbiome [26] showed a sharp increase in phylodiversity after day 161 (Fig. 3), the same date that the subject began consuming baby formula. This suggests that baby formula changed the phylogenetic colonization patterns of the developing infant gut. We analyzed this data set as two separate time-series, one before formula use and one during, and both had negative *D* estimates, with the pre-formula *D* estimate being lower (Fig. 4). While the pre-formula data set was significantly underdispersed (*P* = 0.007), the formula data set was not significantly different from the neutral model, although this result is marginal (*P* = 0.107). When the “individual null” was used instead, both were significantly underdispersed. Infant gut data from Finnish infants [21] were sampled at a much lower temporal resolution, and as such were not split between formula use. 31 out of 33 individuals analyzed exhibited significant underdispersion, and the other two were not significantly different from the neutral model. Both non-significant individuals were from the group treated with heavy antibiotics, but even so, no significant difference in *D* values was detected between antibiotics and control groups (Fig. S4). However, when the “individual null” was used, all subjects exhibited significant underdispersion. Estimates of *D* did not significantly correlate with the number of zOTUs in a data set, the total phylodiversity of the data set, the initial phylodiversity of the data set, or the number of samples in a data set (Fig. S5).

**Fig. 3.**
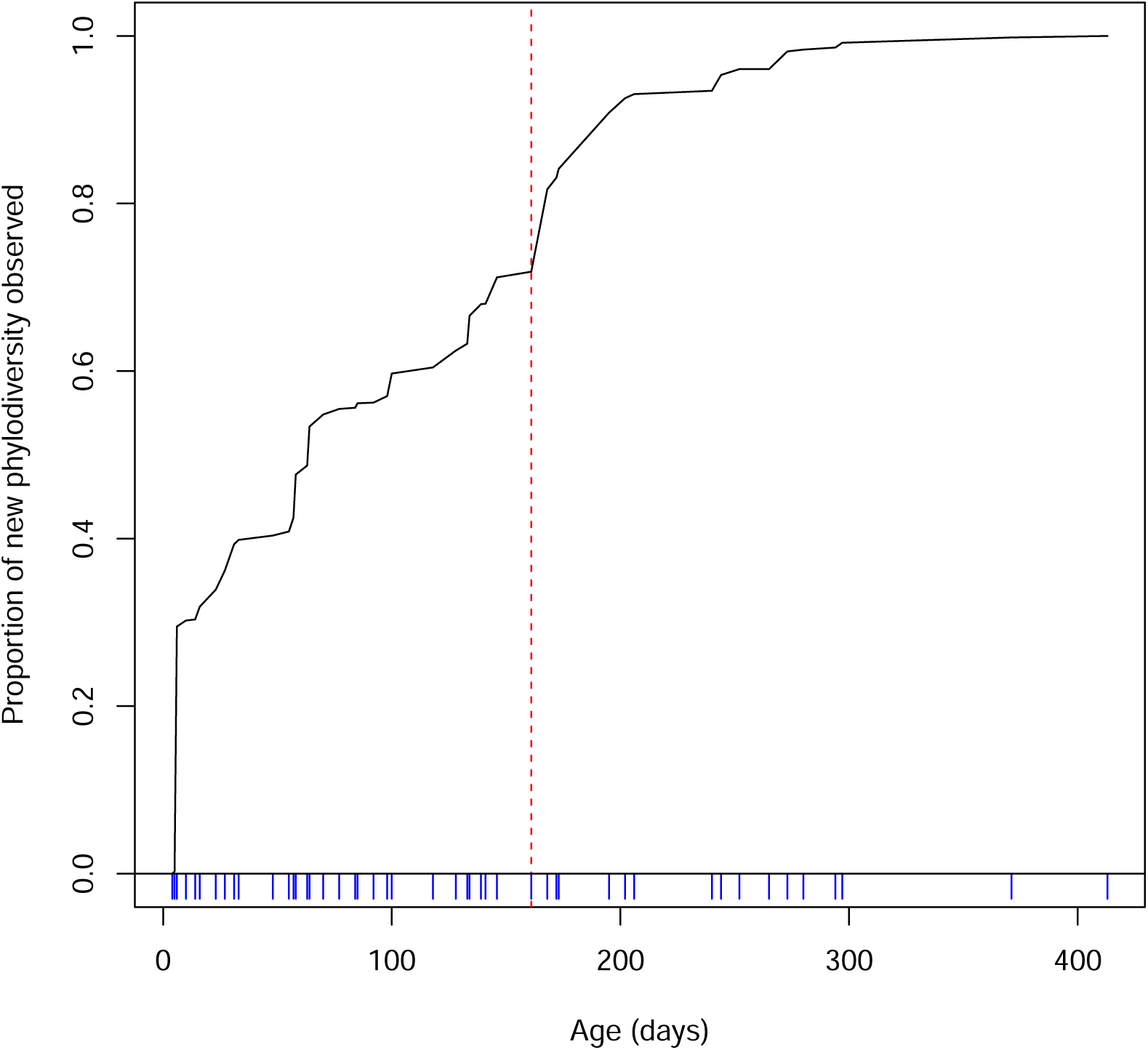
Empirical phylodiversity accumulation in the infant gut microbiome [26]. Phylodiversity increases sharply after day 161 of the infant’s life, then plateaus. This timing coincides with the day the subject began consuming baby formula. The times of sampling points are shown as vertical blue lines below the X-axis.

**Fig. 4.**
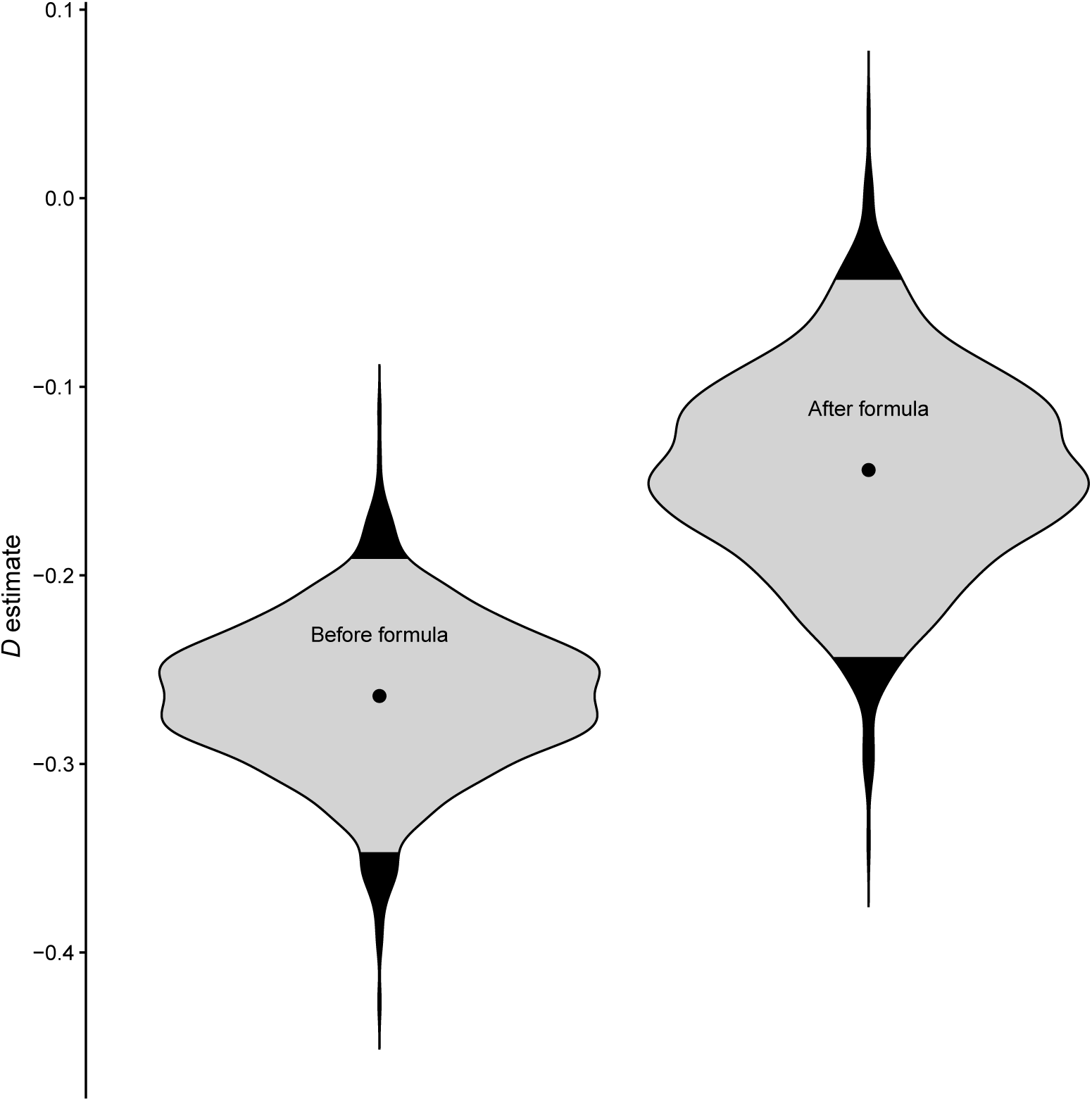
Dispersion parameter (*D*) estimates in the infant gut, pre-formula and during formula use. Formula use began on day 161, thus the first 160 days of the subject’s life were analyzed separately. Community assembly was significantly underdispersed in the pre-formula data set, but was not significantly different from the neutral model during formula use (*P* = 0.107).

## Discussion

Any organism of interest in a human microbiome data set, from the pathogenic to the probiotic, will at some point be recruited for the first time, and the order in which these organisms are detected in the community is determined by community assembly processes [1]. Predicting which lineages of organisms can be recruited into a given environment has far-reaching implications for ecosystem remediation and management, especially in microbial communities where the medical and ecological importances of many microbes are still largely unknown [29; 30]. Identifying conditions under which assembly mechanisms change, or under which non-neutral assembly is particular strong, may facilitate microbial community rehabilitation by understanding when and how microbial communities can be colonized by close/distant relatives. We found that assembly during primary succession of the infant gut (Fig. 4, Fig. S4) and during turnover of the microbial communities on the adult palms and gut (Fig. 2) follows a predictable pattern: new species are more likely to be recruited if a close relative has been recruited previously.

The generally “nepotistic” pattern we observed supports our underdispersion hypothesis, which follows Darwin’s pre-adaptation hypothesis [11] and more recent ecological theory as well [18; 19]. Much work in phylogenetic community ecology posits that competition tends to be strongest among closely-related species due to phylogenetic niche conservatism [8], so many closely-related species are able to coexist in a community, competition must not be an important factor structuring that community [20]. However, strong competition between distantly related species may actually cause groups of phylogenetically similar species to coexist, especially when immigration is high [18; 19; 31]. This type of competition is perhaps better conceptualized as selection (*i.e.* environmental filtering) instead [19], especially since studies showing evidence for competitive exclusion in microbial communities focus on competition between closely-related species [13; 2].

Our model investigates the extent to which newly recruited species are likely to be similar to previously recruited close relatives, but “previously recruited” may include a significant time span. Thus, the observation of underdispersion may not reflect a lack of importance of competition between close relatives *per se*. However, asking whether new species recruitment is likely after recruitment of a close relative has relevance; for instance in human microbiome systems it may be beneficial to understand if a pathogen’s probability of recruitment may be higher if a conspecific strain was previously observed [14; 16]. Approaches that consider only recent community membership may more directly inform hypotheses regarding direct competition between close relatives, or regarding more recent recruitment of close relatives. For this reason, we included a sliding-window analysis of 5-day intervals for a subset of intensively-sampled data, and showed significant underdispersion in a majority of windows analyzed (Fig. S3). This type of analysis can satisfy the issue of recency when using our model, but only when data collection is sufficiently frequent.

Regardless, non-neutral patterns of phylogenetic community structure have been interpreted to mean that traits are under ecological selection [32; 20; 33; 34]. If traits are not driving community assembly [35] or if the traits driving community assembly are largely horizontally transferred between taxa independent of their relatedness (as estimated by a 16S rDNA phylogeny), we would expect no phylogenetic signature, and a *D* estimate that is not significantly different from 0 (the neutral model). Instead, we observed a very strong and significant phylogenetic signal in species recruitment order for almost all data sets we analyzed. However, if selection on traits is driving this pattern, selection itself may not occur within the host environment. An alternative explanation for the underdispersion we observed is that selection is external to the host environment (*i.e.* selection occurs within the neighboring species pool from which emigration occurs), causing change in the community entering the host to already be underdispersed. Similarly, phylogenetic dispersion of community structure has been unable to distinguish between selection and differences in migration rates [36], so a pre-underdispersed community entering the host is a plausible mechanism for phylogenetic under-dispersion of species recruitment. But selection of microbial communities within the host has been shown by multiple studies [37; 38; 39], so it is our opinion that selection within the host is a more likely scenario.

A similar point is that species compositions of these data sets do not reflect the actual species pool from which they originate, which is likely not static over time. Our model draws species from the pool of all species observed over the course of a time-series, and with that model we ask whether the order in which those species were recruited is consistent with under- or overdispersion hypotheses. It is for this reason that we discuss our model in the context of the order of recruitment, regardless of whether selection took place in the host or was external as described above. A null model that accounts for a dynamic species pool that is external to the host could be used with the type of analysis we present here, in order to understand if the processes underlying over-/underdispersion take place in the host or in the host’s environment. Our “meta vs. self” comparison (Fig. S2) uses the combined habitat-specific microbiomes of two co-habitating individuals as a broader species pool, and we found even stronger underdispersion when empirical “self” patterns were compared to the broader “meta” species pool, suggesting strong selection within the host environment. Still, this broader species pool is not a substitute for a thorough inventory of the real species pool, which would be required to ascertain whether over/underdispersion originated in the host or elsewhere. As such, future studies may wish to intensively sample a subject’s environment in order to better catalog the species pool, and perhaps use qPCR to obtain environmental concentrations of species within the species pool [40].

As to why no data sets analyzed showed significant phylogenetic overdispersion (*D* > 0), we are not certain. At the beginning of development of this model, we expected microbial communities in the human microbiome to follow the overdispersion hypothesis, partly from microbiome studies suggesting competition among closely-related bacteria is an important factor in human gut microbial community assembly [41; 42; 1], and also because of work in experimental microcosms [13]. However, the human microbiome environments analyzed here are environments that undergo constant physical disturbance, unlike aqueous microcosms. Palm communities are physically disturbed with every use of the hands, and by the sampling procedure itself. Gut (fecal) communities are also disturbed constantly by the movement of feces through the gut. It may be possible that continuous disturbance allows for underdispersion via constant re-assembly of communities. In this case, niches may be filled by random “winners” after each disturbance, as in a competitive lottery scenario [4]. These “winners” would still need to be pre-adapted to their environment, so they would be more likely to be closely related to previous “winners”, as in our findings. Similarly, environments with fluctuating resource profiles may result in clusters of organisms occupying the same niche [43]. The data sets we used are also somewhat limited in terms of phylogenetic resolution, as short reads of the 16S marker gene are insufficient to detect strain-level variation [44; 42; 15]. Thus, competitive exclusion could occur at the extreme tips of the bacterial phylogenetic tree, and this would not be detectable using 16S rDNA data. Even so, broader patterns of underdispersion at phylogenetic depths accessible with 16S data could still result in significantly underdispersed model fits.

A strength of our model is that it estimates values of *D* that can be compared among data sets (Fig. 2) or potentially across time (Fig. 4, Fig. S3) in order to learn how differences between data sets impact community assembly. We found that gut and palm communities were almost universally underdispersed (Fig. 2, Fig. 4, Fig. S4), and that the D value for a community appears to be a function of body site (Fig. 2). Although this result is only shown across two subjects, the parallel patterns between the male and female subject are striking, in that fecal communities are the most strongly underdispersed (lowest *D*), palm communities are similar to each other, and tongue communities had the highest *D* estimates. Similarly, comparing *D* before and after an event can be used within an experimental framework to see how that event may affect community assembly. Our analysis of infant gut microbiome data [26] before and during the use of baby formula (Fig. 4) showed that while the pre-formula community was significantly underdispersed, community assembly during formula consumption was more neutral. While the post-formula trend was not significantly different from the neutral model, this finding was marginal (*P* = 0.107).

In addition to showing that our model can be a useful tool for future studies, our findings also hint that phylogenetic underdispersion may be a common trend for the human gut microbiome, although demonstrating a general trend would require analysis of more than the 36 individuals we analyzed. Indeed, recent research has shown that for fecal transplants, donor strains are able to integrate into the recipient’s gut community when a conspecific strain is already present, but novel donor strains are unlikely to successfully integrate into the recipient [14]. Congeneric bacteria have also been shown to be predictors of each others’ recruitment in the mouse gut microbiome, both for pathogens and commensals [16]. Different body sites - as we saw with the skin – may have qualitatively similar patterns of underdispersion, yet quantitatively different magnitudes of this effect. Thus the efficacy of an engineered probiotic based on similarity to organisms already present in the community for which it was engineered may largely depend on the body site for which it’s intended, although again more exhaustive study is needed. To facilitate further discovery both in the human microbiome and in other environments, we have made our R code and a tutorial available on GitHub: https://github.com/darcyj/pd_model.

## Acknowlegements

The authors thank D.R. Nemergut for her help and support, and also thank P. Sommers, E. Gendron, A. Solon, E. Pruesse, A. Armstrong, C. Martin, K. Hazleton, and S. Sauce for many helpful discussions. Funding was provided by an NSF grant for studying microbial community assembly following disturbance (DEB-1258160) and by a NIH NLM Computational Biology training grant (5 T15 LM009451-12).

## Supplementary Material

**Fig. S1.**
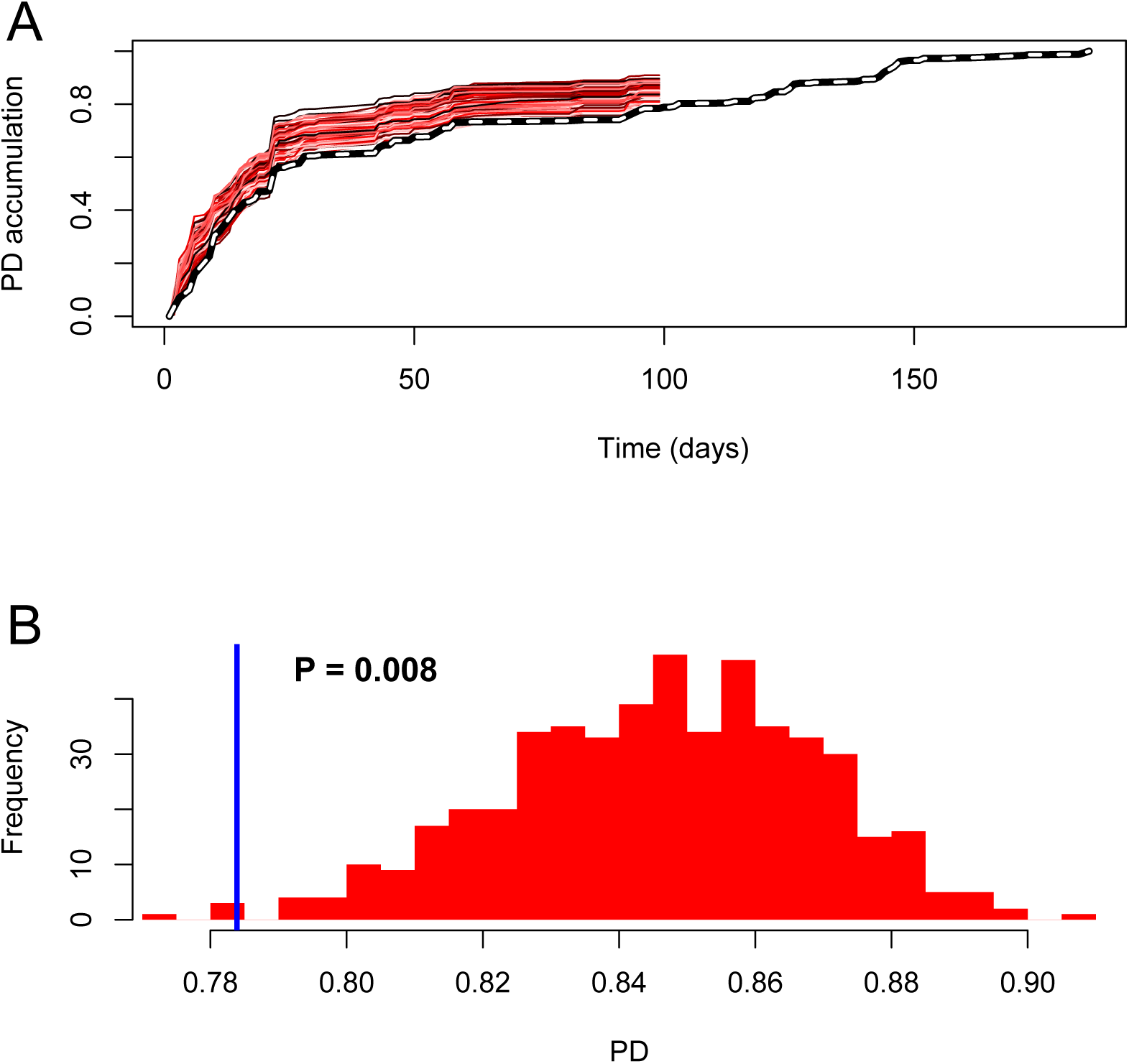
Significance testing for the female feces data set. Plot A shows the empirical phylodiversity accumulation (dashed; same as Fig. 1A) but with neutral model surrogate data sets shown in different shades of red. These are produced by running the neutral model 500 times, to generate a distribution of phylodiversity values under *D* = 0 (Plot B). As with all surrogate data sets, these are run until time *m* (see Parameter Estimation section of Materials and Methods). Empirical phylodiversity at time *m* (blue line) is compared to the distribution of neutral model phylodiversities at time *m* (red histogram), and a *P*-value is calculated as the proportion of neutral phylodiversities more extreme than the empirical value.

**Fig. S2.**
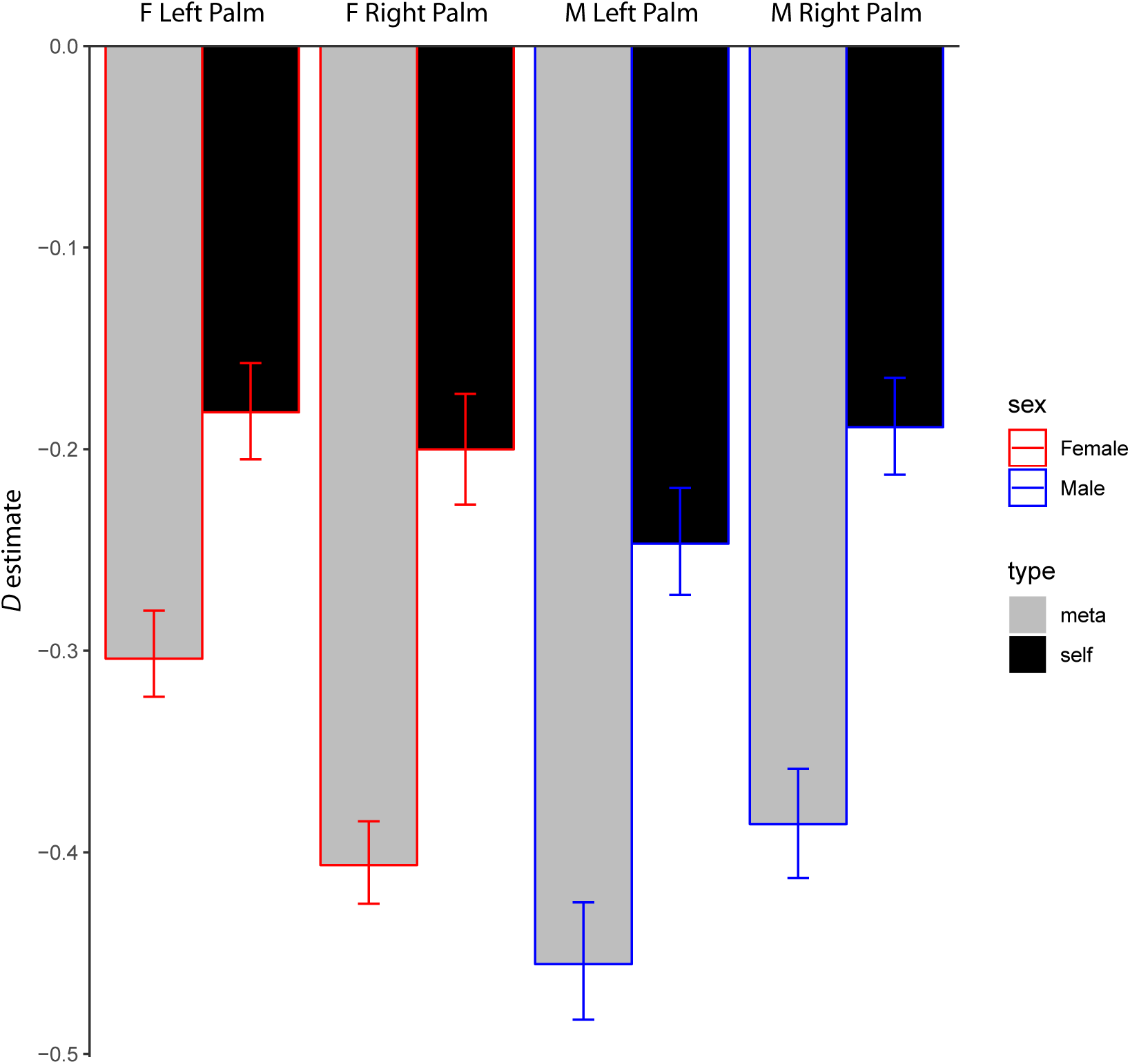
Comparison of “self” vs “meta” model results from palm communities. “Self” (black) models were run identically to Fig. 2), but “meta” (gray) models were run where the species pool for each palm community surrogate data set was composed of all zOTUs observed across all four palm data sets. The difference between the “self” *D* estimate (generated above) and the “meta” *D* estimate (estimated with a metapopulation of zOTUs) is related to the exclusivity of recruitment into the community. In other words, if we were to estimate similar *D* values for both the “meta” and “self” analyses, the inclusion of extra species in the species pool would be of little importance to the model, and we would learn that it would make little difference to community assembly patterns if the species pool really was composed of the “meta” set.

**Fig. S3.**
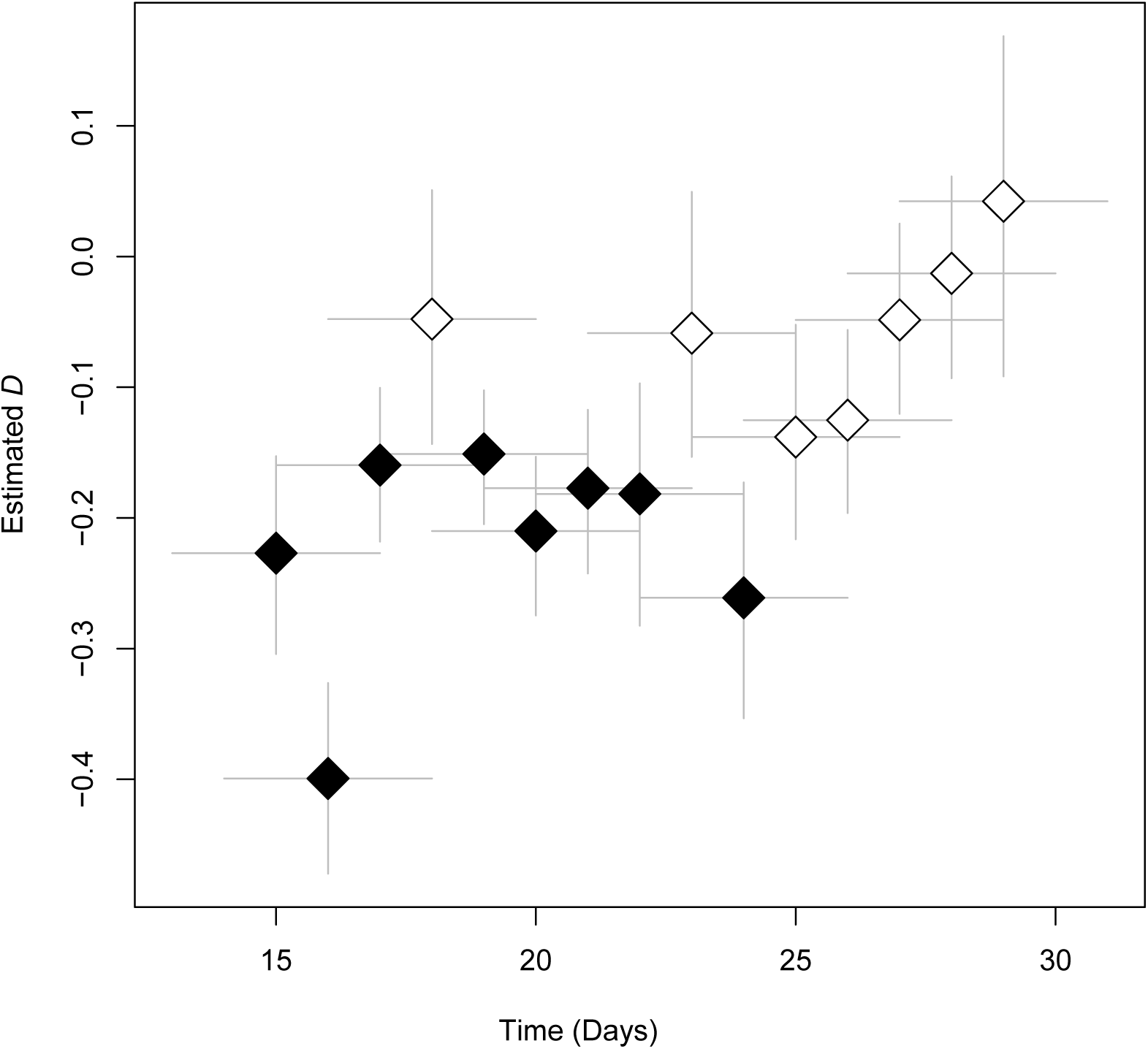
Sliding window analysis of male right palm data over 19 consecutive samples. We ran our model on each window of 5 continuous days (15 windows), in order to see how *D* varied over time. We only conducted this analysis for the section of samples that were sampled every day, so that comparisons between windows would not be confounded by window size. This analysis was done to demonstrate a potential use case for our model, and not to test any specific hypothesis. Filled shapes represent windows that were significantly different than the neutral model. Vertical bars represent 95% confidence intervals for *D* estimate, and horizontal bars represent window size.

**Fig. S4.**
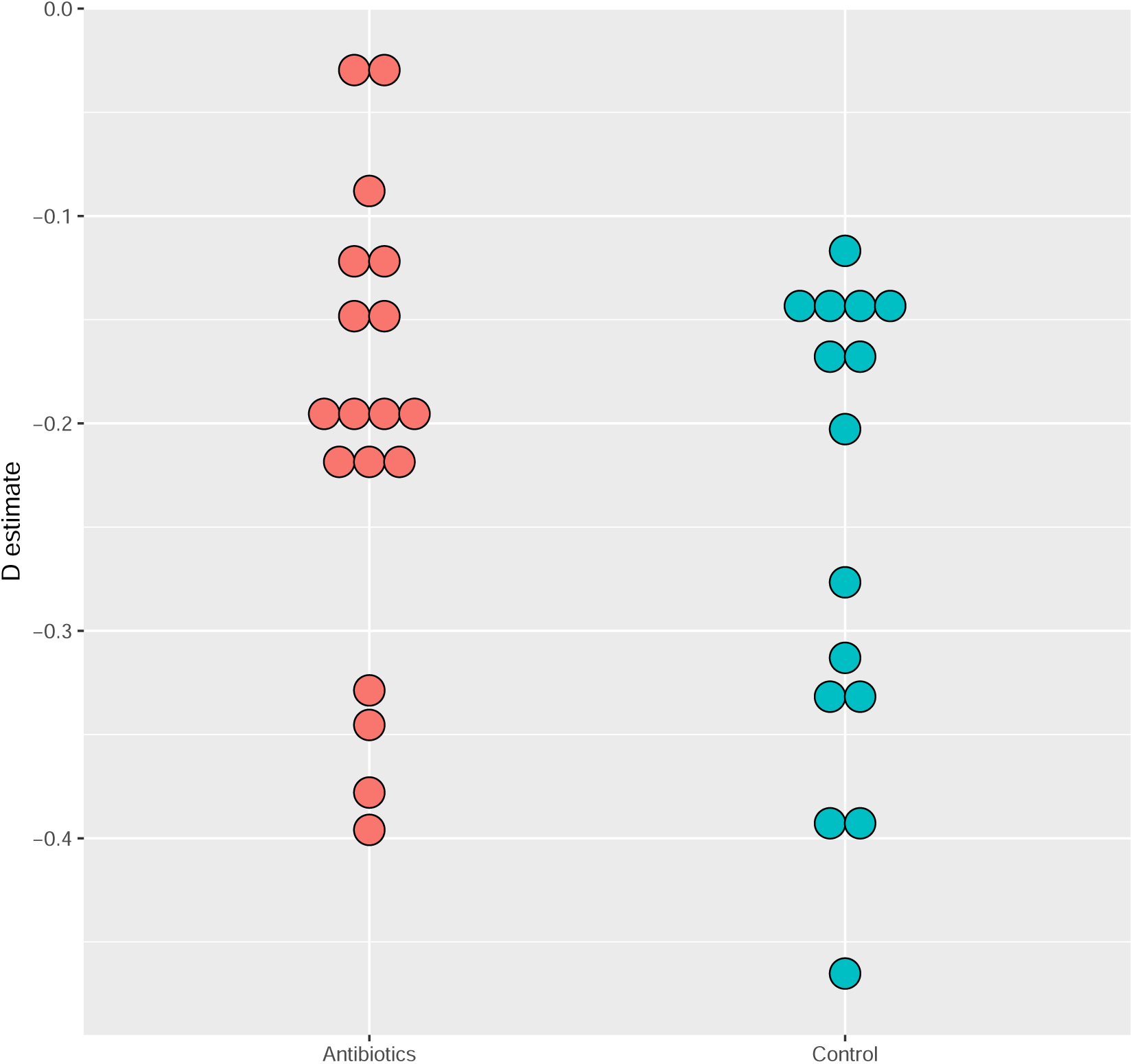
*D* estimates of Finnish infant data sets. All but two subjects exhibited significant phylogenetic underdispersion. The two subjects that were not significantly different from the neutral model were both in the antibiotics cohort, which is comprised of infants that were treated with frequent antibiotics, almost all for ear infections. There was no significant difference between *D* values for the two groups.

**Fig. S5.**
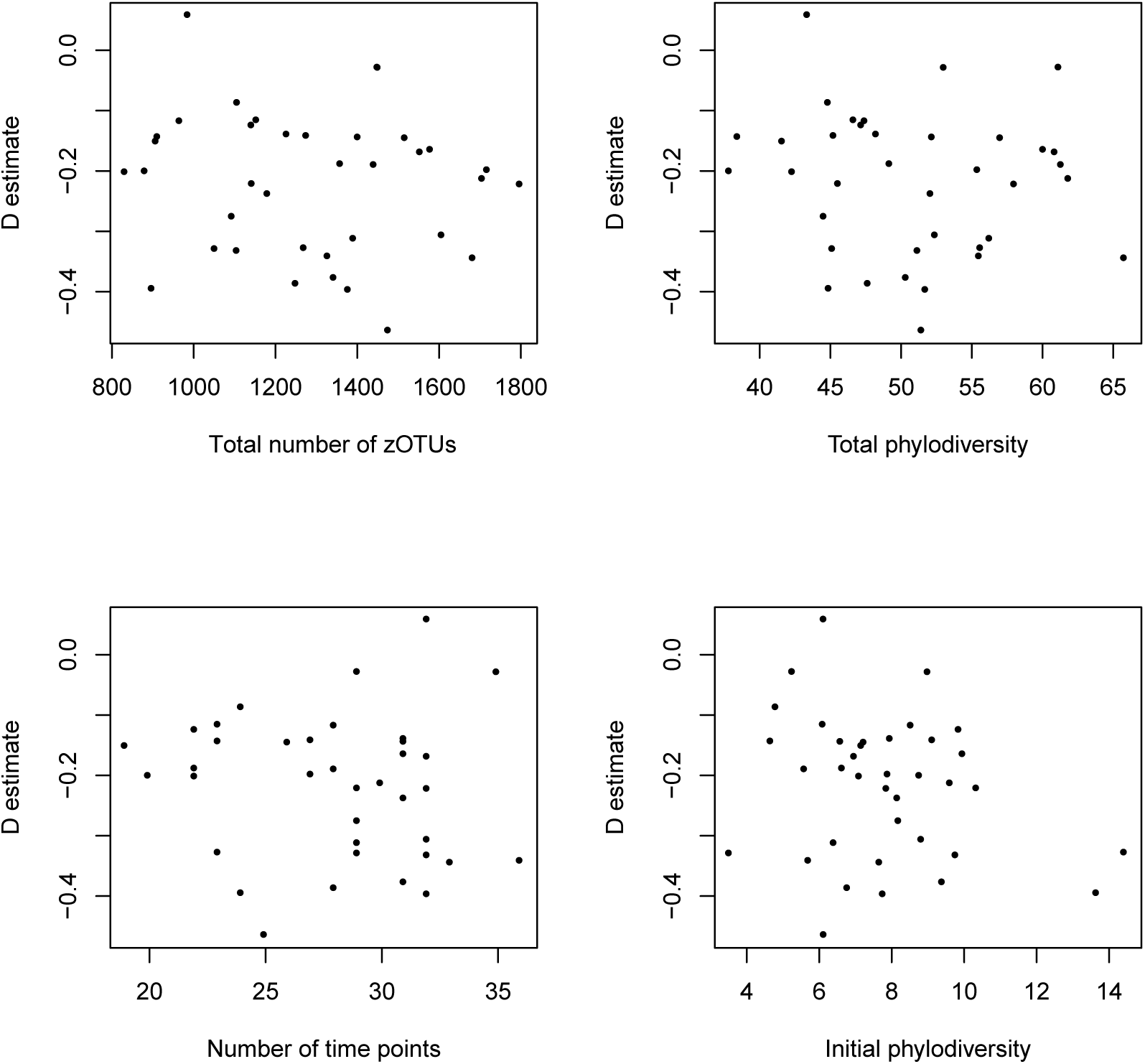
Relationship of *D* estimate to total zOTU richness, total phylodiversity, number of timepoints sampled, and initial phylodiversity (of first sample) for Finnish infant data. No statistically significant correlation was detected in any of these four analyses.

